# Smells allow for tapping into episodic-like memory of minipigs

**DOI:** 10.1101/2025.07.11.664385

**Authors:** Mariam Hillmann, Edna Hillmann, Lorenz Gygax Hillmann

## Abstract

Episodic-like memory is considered as one of the most advanced cognitive capacities and could therefore have major implications on how animals with this capacity are treated in respect to their welfare. Yet, episodic-like memory has only been substantiated in a few vertebrate species. These species are quite diverse, originate from several taxa, and the question is raised whether episodic-like memory is indeed a rather specific capacity or whether this capacity turns out to be more common when further species are investigated. In general, intensively foraging animals that rely on some replenishing resources are promising species to investigate episodic-like memory. Here, we conducted a pilot study on minipigs (*Sus scrofa f. domestica*), replicating an approach used in dogs. Both, dogs and pigs are macrosmate species, making the comparison of their memory of olfactory stimuli promising. The minipigs learnt to differentiate the sequence of encountered location-smell combinations as easily as dogs. This makes minipigs a promising species and the use of smells a promising approach to investigate episodic-like memory in more detail. This study also hints towards a cognitive capacity of pigs, which raises further concerns regarding intensive and barren housing conditions.

## 1 Introduction

Episodic memory is the human capacity to recall a specific past personal event. Tulving (2005) argued that episodic memory is based on mentally traveling back in time to subjectively remember a specific event (autonoetic consciousness). Yet, direct access to episodic memory is limited in non-human animals due to the lack of verbal interaction. Accordingly, episodic-like memory has often been investigated by the capacity of remembering what happened where and when in non-human animals (Hampton & Schwartz 2004, Breeden et al. 2016, Madan 2020, Huston & Chao 2023). Moreover, this capacity has been viewed as one of the most advanced cognitive capabilities in animals (Pladevall et al. 2020, but see: Allan & Fortin 2013), which may have important welfare implications in respect to how animals with this capability are treated. Not surprisingly given the high level of cognition, there is support for episodic-like memories in only a few animal species representing different clades such as rodents (mice: Dere et al. 2005; rats: e.g. Babb & Crystal 2006; long-tailed rats: Yi et al. 2021), dogs (Lo & Roberts 2019, Fugazza et al. 2020), dolphins (Davies et al. 2022), monkeys (Janson 2016, Hoffman et al. 2009), apes (Martin-Ordas et al. 2010, but see: Pladevall et al. 2020), and birds (e.g. Salwiczek et al. 2010). This small selection of species may not solely be due the limitations in cognitive capacities in other species (Allan & Fortin 2013) but may also be due to the selection of model organisms. Pigs are active foragers (Moser et al. 2019) and the capacity to remember what (kind of feed) they found where and when would make foraging clearly more efficient (Fuhrer & Gygax 2017). Accordingly, pigs may present a promising further study species for episodic-like memory. An initial study on (mini)pigs showed some capacity to differentiate stimuli based on what/where/which (Kouwenberg et al. 2009). Here, we conducted another pilot experiment focusing more strongly on the “when” aspect and taking advantage of the olfactory ability of (mini)pigs in an episodic-like memory task. To do so, we followed a protocol developed for dogs (Lo & Roberts, 2019). The main aim of the current study was to assess whether the protocol was applicable to pigs and whether pigs showed some initial capabilities pointing towards episodic-like memory such that further studies using the presented approach will be worthwhile.

## 2 Materials and methods

The experiment was conducted at the Teaching and Research Station for Farm Animal Sciences (Humboldt-Universität zu Berlin, Faculty of Life Sciences, Albrecht Daniel Thaer-Institute of Agricultural and Horticultural Sciences, Berlin, Germany). Twelve female minipigs (7 Göttinger and 5 Aachener) experienced with behavioural tests and aged 5 to 9 years were kept in pairs in neighbouring outdoor pens (10.5 × 2.3 m). Two of these pigs were kept singly due to their low social tolerance but they, too, had visual, olfactory, auditory, and limited physical contact to their neighbours. Pens had a solid ground, a roofed feeding area and huts littered with straw. Pigs were fed twice daily in the early morning and around noon and had ad-libitum access to water and hay.

This experiment was approved by the university’s animal welfare officers and by the responsible veterinary office (LAGeSo Landesamt für Gesundheit und Soziales, Berlin, permit no. StN 00002-23). The veterinary authority assessed the study as an animal experiment that is not regulated under the animal protection law because no pain, suffering, or injury of the tested animals was expected in the course of the study.

Following the procedure of the first of four experiments with dogs, reported by Lo & Roberts (2019), the pigs were asked to differentiate between smells (what) that were provided in a sequence (when) and at specific locations (where). The minipigs had to learn to choose boxes with different smells and were rewarded by small pieces of apple for a correct choice. Testing took place in an experimental pen set-up consisting of the choice area and a start pen with a solid floor as well as a pen with sandy ground where, for the pair-housed minipigs, the pen mate resided during the experiments with visual, olfactory, and auditory contact (Fig. 1). For each experimental run, four target boxes were put in a semi-circle at 100 cm distance from the door of the start pen. The walls of the starting pen were opaque such that animals could not observe the positioning of the target boxes. The door of the start pen was always opened towards the start pen such that all target boxes became approachable simultaneously. Pigs in the start pen and in the partner pen were provided ad-libitum water.

**Fig. 1.**
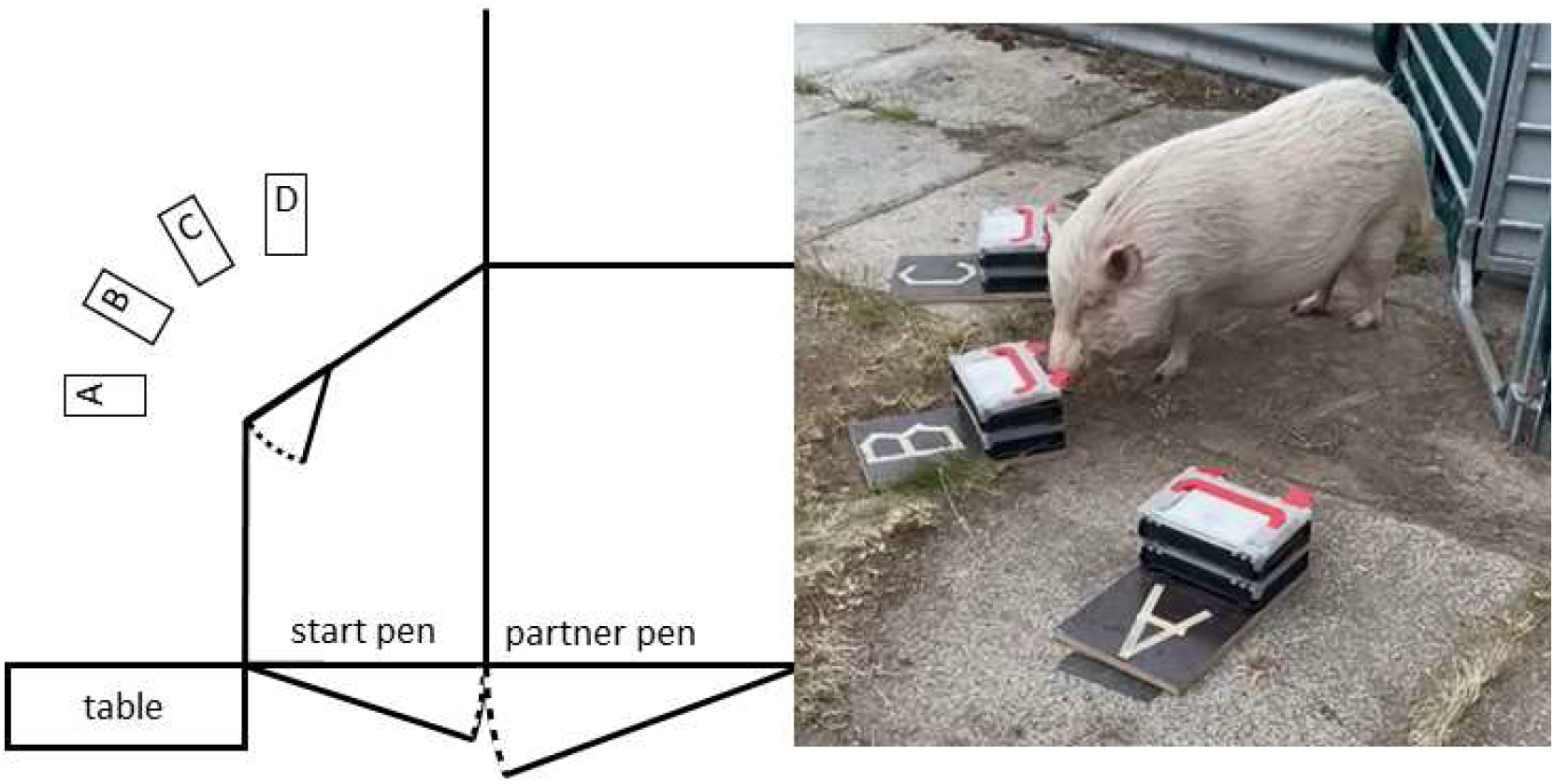
Sketch of the testing area (left) and real testing situation (right). A, B, C, D are the target boxes (see also Fig. 2)

The target boxes consisted of two compartments, one on top of the other (Fig. 2; two boxes of type STRAUSSbox mini, Engelbert Strauss GmbH & Co. KG, Biebergemünd, Germany). The upper contained the apple reward if it was the correct choice. It stayed empty otherwise. The lid to this upper compartment was never locked such that it was accessible in all experimental runs. The lower compartment was perforated and always contained pieces of apple, such that all target boxes smelled of apple. Additionally, it contained a petri-dish with a cotton pad that either had a specific smell (Set of 30 aromatic oils, WTRCSV, available at Amazon Europe, Luxemburg) or some droplets of water applied to it. This corrected for any smell of the cotton pad as well as for the noise that occurred when a petri-dish was put into the lower compartment of the target boxes during preparation or when the box was moved by the animals. This lower compartment was locked and, accordingly, inaccessible to the minipigs.

**Fig. 2.**
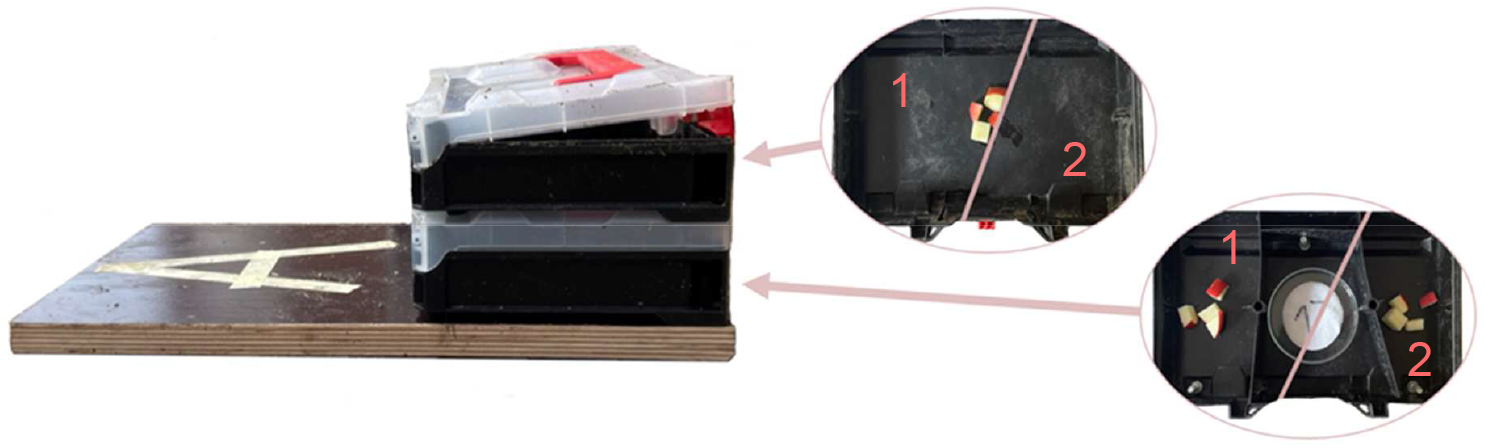
Target box setup. (1) Rewarded: with pieces of apple in the upper box and a numbered smell on a cotton pad in the lower box, (2) un-rewarded: no pieces of apple in the upper box, cotton pad with water in the lower box. For the smell of apple, the lower box always contained some apples.

Boxes were prepared at a table at the edge of the area. Between runs, one experimenter prepared the target boxes and placed them in the choice area. This experimenter always faced the preparation table whenever a minipigs was making a choice. The second experimenter handled the minipig and did not see the preparation and positioning of the target boxes. Therefore, the experimenters could not inadvertently cue the minipigs for the correct target box.

Before the experiment proper, animals were habituated to the test situation and the basic tasks like opening the target boxes and associating a smell with a reward (Table 1). During these habituation sessions, we saw that the experiment lasted quite long, i.e. up to one hour per animal. Moreover, not all minipigs seemed motivated to participate in the task. Some animals were reluctant, other animals explored the choice area but were not interested in the target boxes. Therefore, we selected the six animals (one Aachener and five Göttinger) that were most interested in the target boxes on the habituation days based on the highest number of rewarded boxes that they opened. One additional animal seemed to lose interest in the target boxes later in the experiment and was not further tested after its fifth session.

**Table 1.**
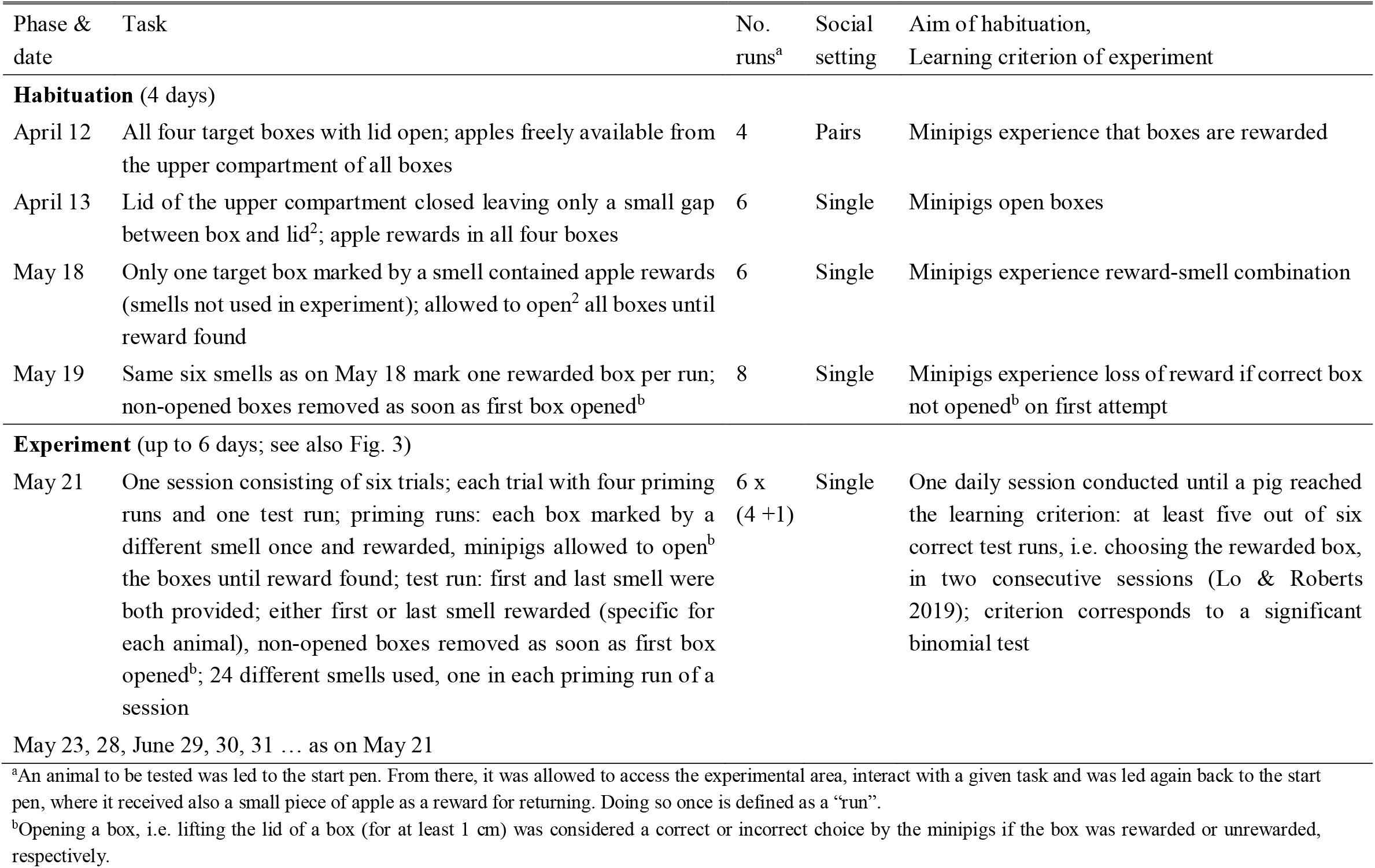
Phases of the experiment, the different tasks per phase, the number of repetitions, social setting, aims of the habituation phases and criterion of experiment.

The steps of the experiment as well as the structure of sessions, trials, and runs are described in detail in Table 1 and illustrated Fig. 3. In all these sessions, the sequence of the smells and the locations in the priming runs were fully randomised using the random number generator in R (Version 4.2.3, R Core Team 2023).

**Fig. 3.**
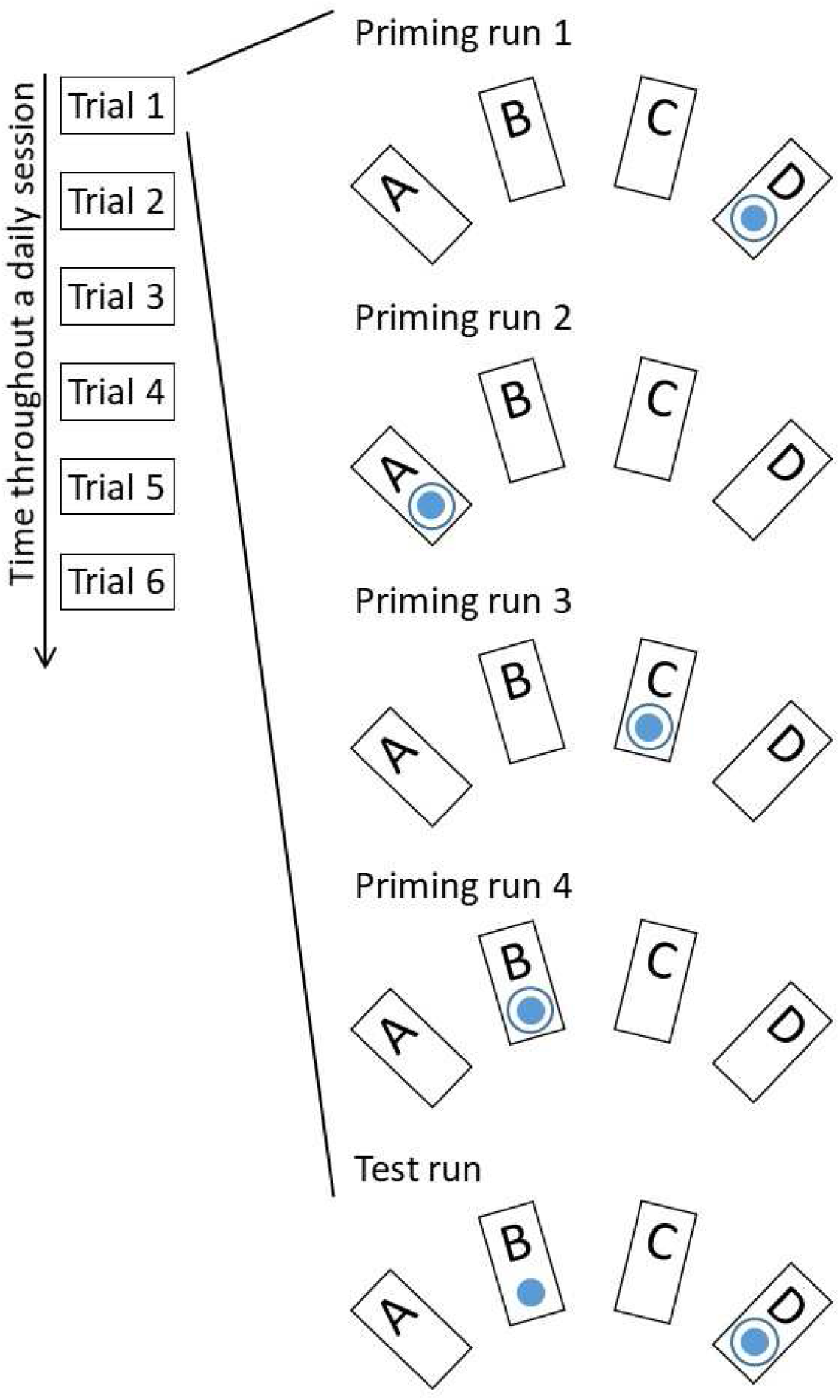
The general temporal course of a daily session with its six trials (on the left) and an exemplary detailed temporal course of one trial including five runs. The first four runs cue one of each box with a smell and a reward. The test runs provides the first and last smell (filled circle) in the locations, where it was presented before, but only rewards either of the two (circle). Here, an example is shown for a pig that is rewarded for the first smell.

## 3 Results and Discussion

The learning curves of the animals increased systematically yet the individual minipigs were quite variable (Fig. 4, top). All the five animals that were continuously motivated to participate in the task did reach the learning criterion in a median of five days (range: 2-6 days). This seemed to be independent of whether animals were trained for the first or last smell-place combination. There was an interesting anecdotal incidence in that one animal reached 6 out of 6 correct test runs in session 3, only 2/6 in session 4 but again 6/6 in sessions 5 and 6. On day 4, the test for this animal was conducted before the noon feeding whereas the animal was tested after the noon feeding in sessions 3, 5, and 6. Seemingly, this animal was not sufficiently able to concentrate when hungry. Overall, we were highly impressed with the patience and concentration that the minipigs showed in this experiment lasting up to an hour per session. Some of minipigs seemed well aware of the procedure in that they went back to the start pen immediately after finding their apple reward during a run expecting their next reward there. Their motivation to participate might have been even higher if the reward was individually tested and assigned. Only a small number of animals was tested here. Yet, the tested animals that showed initial interest in the task were all successful in learning indicating that the task is within the range of cognitive capacities of the species. In conclusion, the minipigs easily learned to differentiate the time order of the smell-location combination showing some necessary if not sufficient capacity for episodic-like memory (Crystal 2021). Overall, we could show that the minipigs learnt the first episodic-like memory task from Lo & Roberts (2019) at least as fast as their dogs who reached criterion in a mean of 5.88 sessions (95% confidence interval [4.57, 7.18]) compared with 4.40 [2.14, 6.66] in our minipigs. It is accordingly promising to continue with further experiments for episodic-like memory in minipigs using the approach tested here. First, additional tests should aim at controlling that minipigs cannot smell the quantity of reward, can solve the task with positions in the temporal sequence other than the first and last, and that they learn more slowly with only information on either location or smell (Lo & Roberts 2019).

**Fig. 4.**
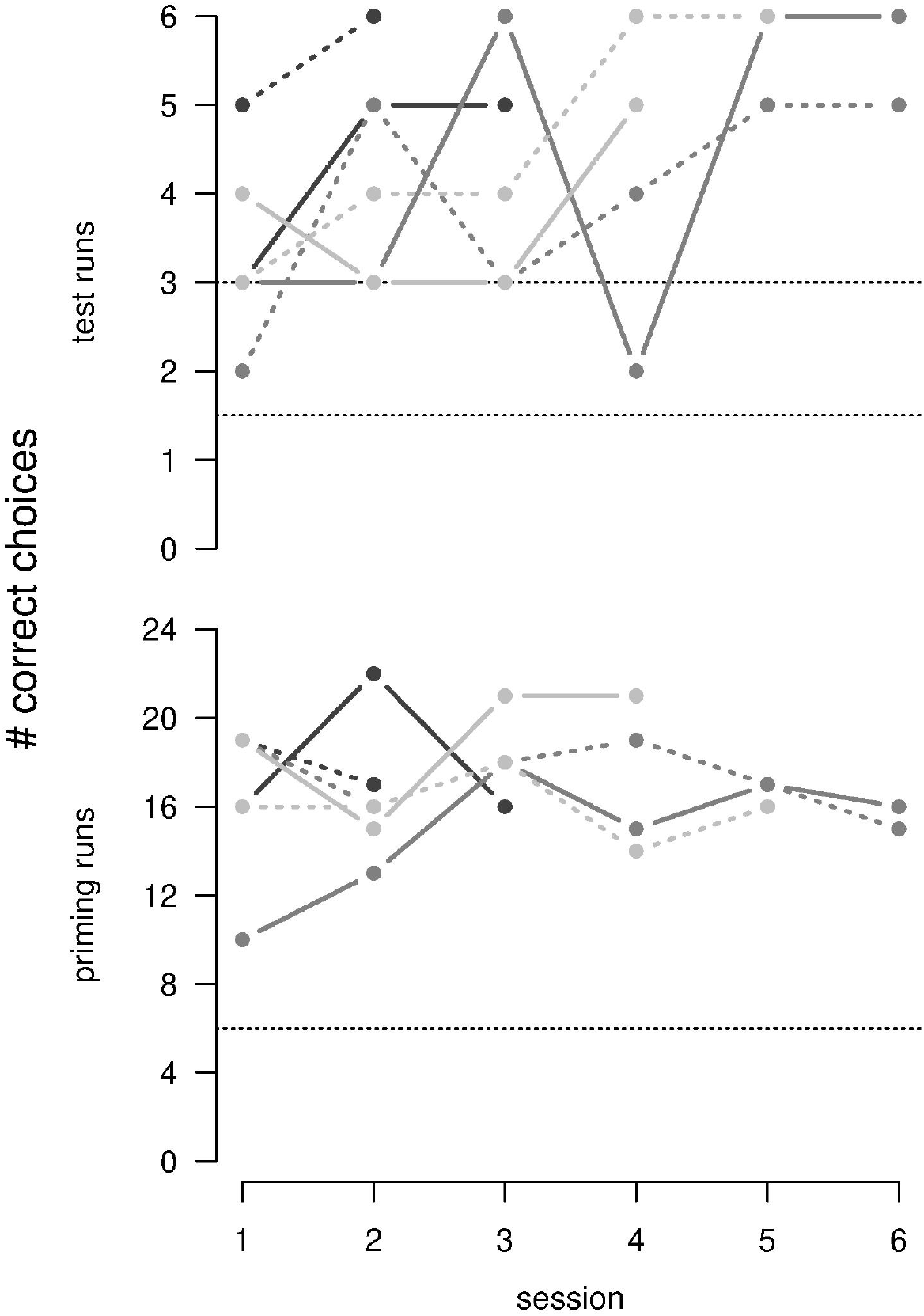
Number of correct choices during the test runs (maximum 6 per session) in the sequence of the sessions (top; horizontal dashed lines: random choices among four boxes or two smells). Number of correct choices (maximum 24 per session) during the priming runs (bottom; horizontal dashed lines: random choice among four boxes). Once the animals reached the learning criterion (two sessions in a row with at least five of six correct choices in the test runs), animals were no longer tested. Each line indicates an individual pig. Dashed line: early smell rewarded, solid line: late smell rewarded

The number of correct initial choices in the priming runs was remarkably constant at around 2/3 of the trials (Fig. 4, bottom). It seems that the pigs tested continuously whether the experimental conditions were still the same (Capaldi & Neath 1995, Jensen 2006) without a detrimental effect on their performance in the test runs.

The cognitive capacity of pigs is moreover relevant in respect to animal welfare. It can be assumed that the common barren housing conditions of pigs, often combined with large amounts of strong smelling gases, restrict welfare all the more if they are so highly cognitively capable as suggested by this experiment.

## Acknowledgements

We thank all the animal caretakers for looking after the minipigs and D. Brucks for commenting on an earlier version of this manuscript.

## Data statement

The datasets generated and analysed during the current study as well as the code used for randomising the conditions of each run can be found in the OSF repository,https://osf.io/y739g.

## References

Allen TA, Fortin NJ (2013) The evolution of episodic memory. PNAS 110:Suppl2:10379–10386. 10.1073/pnas.1301199110

Babb SJ, Crystal JD (2006) Episodic-like Memory in the Rat. Cur Biol 16: 1317–1321. 10.1016/j.cub.2006.05.025

Breeden P, Dere D, Zlomuzica A, Dere E (2016) The mental time travel continuum: On the architecture, capacity, versatility and extension of the mental bridge into the past and future. Rev Neurosci 27:421–434. 10.1515/revneuro-2015-0053

Capaldi EJ, Neath, I (1995) Remembering and forgetting as context discrimination. Learn Mem 2:107–132. 10.1101/lm.2.3-4.107

Crystal JD (2021) Evaluating evidence from animal models of episodic memory. J Exp Psychol: Anim Learn Cogn 47:337–356. 10.1037/xan0000294

Davies JR, Garcia-Pelegrin E, Baciadonna L, Pilenga C, Favaro L, Clayton NS (2022) Episodic-like memory in common bottlenose dolphins. Cur Biol 32:3436-3442.e2. 10.1016/j.cub.2022.06.032

Dere E, Huston JP, De Souza Silva MA (2005) Episodic-like memory in mice: Simultaneous assessment of object, place and temporal order memory. Brain Res Prot: 16:10 – 19. 10.1016/j.brainresprot.2005.08.001

Fugazza C, Pongrácz P, Pogány Á, Lenkei R, Miklósi Á (2020) Mental representation and episodic-like memory of own actions in dogs. Sci Rep 10:1. 10.1038/s41598-020-67302-0

Fuhrer N, Gygax L (2017) From minutes to days – the ability of sows (Sus scrofa) to estimate time intervals. Behav Proc 142:146–155. 10.1016/j.beproc.2017.07.006

Hampton RR, Schwartz BL (2004) Episodic memory in nonhumans: What, and where, is when? Cur Op Neurobiol 14:192–197. 10.1016/j.conb.2004.03.006

Hoffman ML, Beran MJ, Washburn DA (2009) Memory for “What”, “Where”, and “When” Information in Rhesus Monkeys (Macaca mulatta). J Exp Psychol: Anim Behav Proc. 35:143 – 152. 10.1037/a0013295

Huston JP, Chao OY (2023) Probing the nature of episodic memory in rodents. Neurosci Biobehav Rev 144. 10.1016/j.neubiorev.2022.104930

Janson CH (2016) tCapuchins, space, time and memory: An experimental test of what-where-when memory in wild monkeys. Proc Royal Soc B: Biol Sci 283:1840. 10.1098/rspb.2016.1432

Jensen R (2006) tBehaviorism, latent learning, and cognitive maps: Needed revisions in introductory psychology textbooks. Behav Anal 29:187–209.

Kouwenberg A-L, Walsh CJ, Morgan BE, Martin GM (2009) Episodic-like memory in crossbred Yucatan minipigs (Sus scrofa). Appl Anim Behav Sci 117:165–172. 10.1016/j.applanim.2009.01.005

Lo KH, Roberts WA (2019). Dogs (Canis familiaris) use odor cues to show episodic-like memory for what, where, and when. J Comp Psychol 133:428–441. 10.1037/com0000174

Madan CR (2020) Rethinking the definition of episodic memory. Can J Exp Psychol 74:183– 192. 10.1037/cep0000229

Martin-Ordas G, Haun D, Colmenares F, Call J (2010) Keeping track of time: Evidence for episodic-like memory in great apes. Anim Cogn 13:331–340. 10.1007/s10071-009-0282-4

Moser J, Burla J., Gygax L (2019) Executing specific foraging behaviours does not represent a general goal state of foraging in dry sows (Sus scrofa). Behav Proc 164:115–122. 10.1016/j.beproc.2019.05.005

Pladevall J, Mendes N, Riba D, Llorente M, Amici F (2020) No evidence of what-where-when memory in great apes (Pan troglodytes, Pan paniscus, Pongo abelii, and Gorilla gorilla). J CompPsychol 134:252–261. 10.1037/com0000215

R Core Team (2023) R: A Language and Environment for Statistical. Computing. R Foundation for Statistical Computing, Vienna, Austria. https://www.R-project.org/

Salwiczek LH, Watanabe A, Clayton NS (2010) Ten years of research into avian models of episodic-like memory and its implications for developmental and comparative cognition. Behav Brain Res 215:221–234. 10.1016/j.bbr.2010.06.011

Tulving E (2005). Episodic memory and autonoesis: Uniquely human? In Terrace HS & Metcalfe J (Eds.), The missing link in cognition (pp. 4–56). New York, NY: Oxford University Press

Yi X, Yi S, Deng Y, Wang M, Ju M (2021) High-valued seeds are remembered better: evidence for item-based spatial memory of scatter-hoarding rodents. Anim Behav 175:1–6. 10.1016/j.anbehav.2021.02.009

Zentall TR (2013) Animals represent the past and the future. Evol Psychol 11: 573–590. 10.1177/147470491301100307

